# The native structure of the full-length, assembled influenza A virus matrix protein, M1

**DOI:** 10.1101/2020.06.24.168567

**Authors:** Julia Peukes, Xiaoli Xiong, Simon Erlendsson, Kun Qu, William Wan, Leslie J. Calder, Oliver Schraidt, Susann Kummer, Stefan M. V. Freund, Hans-Georg Kräusslich, John A. G. Briggs

## Abstract

Influenza A virus causes millions of severe illnesses during annual epidemics. The most abundant protein in influenza virions is the matrix protein M1 that mediates virus assembly by forming an endoskeleton beneath the virus membrane. The structure of full-length M1, and how it oligomerizes to mediate assembly of virions, is unknown. Here we have determined the complete structure of assembled M1 within intact virus particles, as well as the structure of M1 oligomers reconstituted in vitro. We found that the C-terminal domain of M1 is disordered in solution, but can fold and bind in trans to the N-terminal domain of another M1 monomer, thus polymerising M1 into linear strands which coat the interior surface of the assembling virion membrane. In the M1 polymer, five histidine residues, contributed by three different M1 monomers, form a cluster that can serve as the pH-sensitive disassembly switch after entry into a target cell. These structures therefore provide mechanisms for influenza virus assembly and disassembly.

---

Influenza A virus (IAV) is a negative sense single stranded RNA virus that causes 3-5 million cases of severe illness during annual epidemics. The 8 viral genome segments, their encapsidating nucleoproteins (NP) and the viral polymerase complexes are protected by a lipid envelope derived from the host cell membrane. The exterior of the envelope is densely decorated by the membrane-anchored glycoproteins haemagglutinin (HA) and neuraminidase (NA). A few copies of the ion channel M2 are embedded in the virus membrane, while the matrix protein 1 (M1) is tightly associated with the inner surface of the viral membrane ^1^.

Influenza virions are pleomorphic, producing filamentous and spherical virions ^2,3^. M1 plays a critical role during virus assembly. Co-expression of M1 together with either of the viral glycoproteins, HA and NA, is sufficient to form filamentous protrusions, which in the presence of matrix protein 2 (M2) are released as particles that closely resemble virions ^4^. M1 interacts with viral ribonucleoproteins (vRNPs) ^5,6^ and the cytoplasmic tails of the viral glycoproteins HA and NA ^1,7,8^ to promote their incorporation into virions. After uptake into a target cell, pH-induced structural changes in M1 are thought to contribute to virus entry and disassembly ^9^.

The 252-residue long, 28 kDa M1 protein consists of an N- and C-terminal domain (NTD and CTD). The structure of the NTD has been determined by crystallography, revealing a globular fold of 9 alpha-helices and a highly charged surface ^10–12^. There are no structures available for the CTD of M1. A structure of the full-length matrix protein is available from an orthomyxovirus Infectious Salmon Anemia Virus (ISAV), however substantial divergence (18% sequence identity) between ISAV and IAV matrix proteins makes it unclear how relevant the ISAV matrix protein structure is to IAV M1 ^13^. Classical electron microscopy suggests that M1 forms a helical arrangement of fibers in both filamentous ^14,15^ and spherical virions ^16^, but how M1 oligomerizes and is arranged to form the viral endoskeleton is unknown. The lack of structural data currently hampers understanding of how M1 functions to assemble and disassemble virions. In this study, we combined cryo electron tomography (cryoET) of M1 in situ within IAV virions with cryo electron microscopy (cryoEM) of an in vitro reconstituted helical array of M1 to determine the structure of full-length, oligomerized M1.

To avoid any artefacts associated with purification ^17^ we performed cryoET on influenza A/Hong Kong/1/1968 (H3N2) (HK68) virions directly budding from infected cells ^18,19^. We observe filamentous virus particles radiating from infected cells as well as isolated filamentous virions (**Fig. 1a**). Within the reconstructed tomograms, a dense glycoprotein layer can be seen on the outer surface of the virus envelope, and a thin M1 layer directly underneath the viral membrane (**Fig. 1b**).

**Figure 1:**
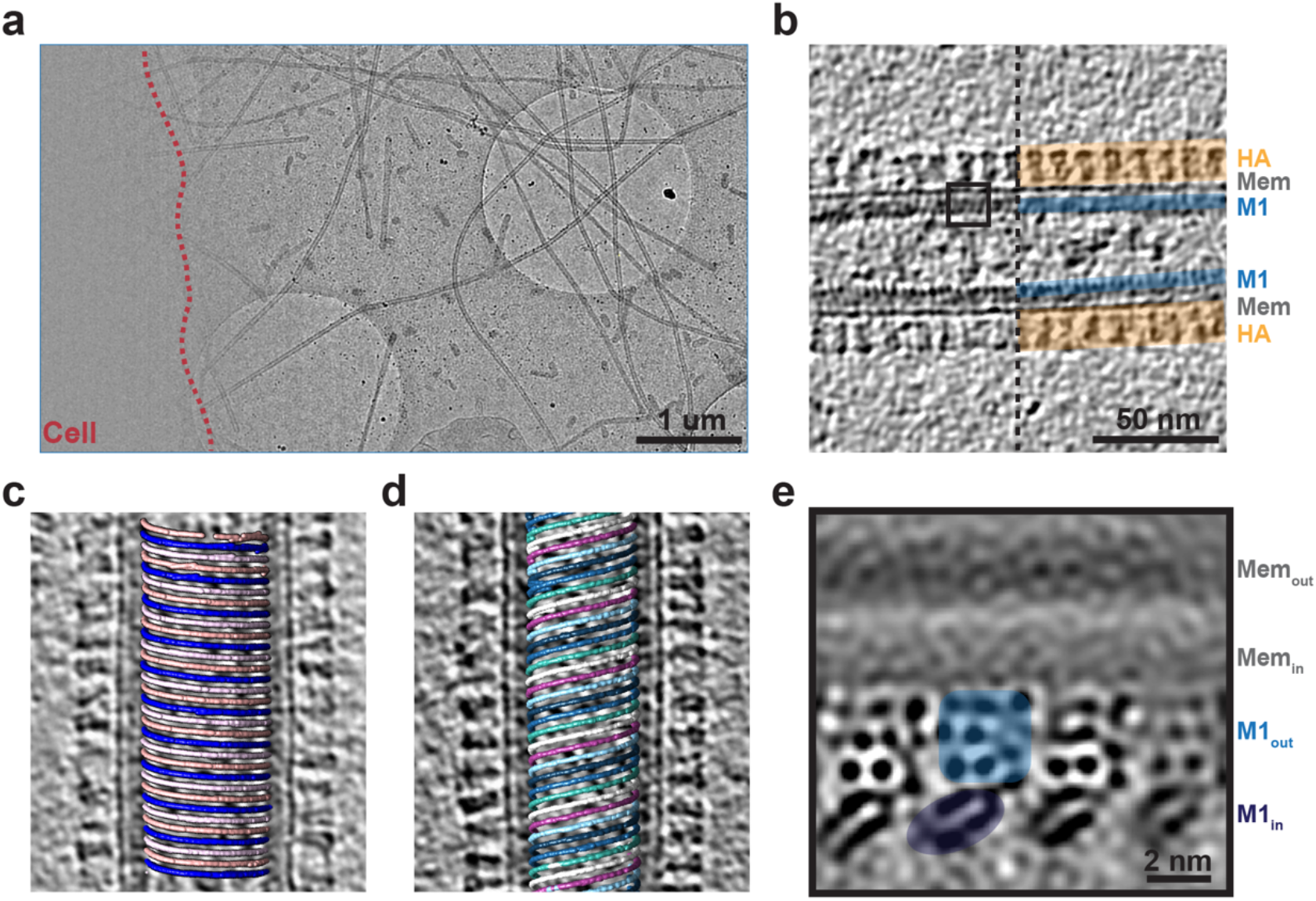
CryoET of influenza A/HK68 virions. **a)** 2D cryoEM image of influenza A/HK68 virions surrounding an infected cell (outlined in red). **b)** Central slice through a tomogram of an individual influenza A/HK68 virion. The surface HA layer (orange), the membrane, and the M1 matrix layer (blue) are marked. **c)-d)** M1 assembles parallel helical strands, visualized by marking the positions of aligned M1 subtomograms within the original tomogram volumes. **c)** a left-handed helix with 3 parallel M1 strands. **d)** a right handed helix with 6 parallel M1 strands. **e)** Orthoslice through four parallel strands of the M1 structure determined by subtomogram averaging. The membrane bilayer and the outer and the inner lobes of the M1 reconstruction are marked. Dark densities correspond to slices through protein alpha-helices. The size and orientation of the view correspond to the black square in b).

We extracted subtomograms containing the M1 density from reconstructed tomograms and performed reference-free subtomogram averaging for each virion individually. By displaying the aligned positions and orientations in 3D, we found that M1 forms tightly packed, linear polymers that assemble a helical array (**Fig 1c,d**, **Fig. 2a**). The spacing between strands, ~ 3.6 nm, is similar to those observed in classical EM experiments ^14–16,20^. The diameter of the virions, and the number of parallel polymer strands, both vary (1-6 strands at a radius from 18 – 29 nm, **Extended data Fig. 1**). M1 subtomograms from virions with three strands were combined into a larger data set for subsequent iterations. The resulting reconstruction of M1 in situ shows the inner and outer leaflet of the membrane bilayer with a two-lobed protein layer bound tightly to its inner surface (**Fig. 1e**, **Extended data Fig. 2a-c**).

**Figure 2:**
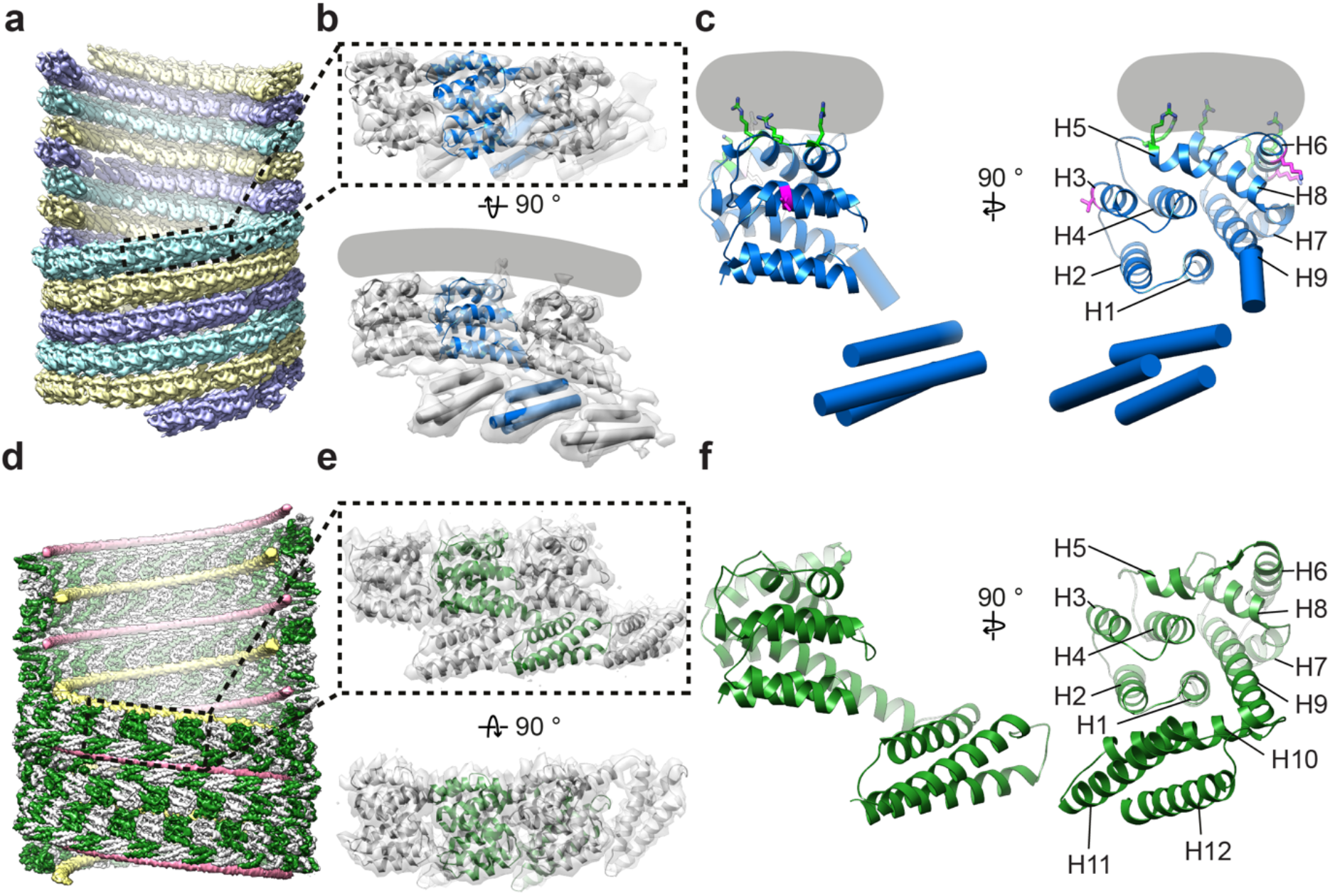
Structure and assembly of influenza M1. **a)** Arrangement of M1 within influenza A/HK68 HANAM1M2 VLPs visualized by placing the M1 monomer structure at orientations and positions determined by subtomogram averaging. Three parallel strands which are coloured differently. Some monomers have been removed to reveal the inside of the filament. **b)** The structure of three neighboring M1 monomers in a strand determined by subtomogram averaging (grey surface), fitted with a crystal structure of the M1 NTD (PDB:1ea3) and a secondary-structure model for the M1 CTD. The grey line indicates the position of the membrane. **c)** The model of an individual M1 monomer. Helix (H) numbers are indicated. Positively charged residues making membrane interactions are green (residues 76 and 78 in H5, 101 and 104 in H6, 134 in H8). Residues at the inter-strand interface where mutation alters virion morphology are magenta (residue 41 in H3, 95 and 102 in H6). **d)** Arrangement of M1 (green and white) within helical assemblies of M1 assembly in vitro. Nucleic acid strands are shown in yellow and pink **e)** Three neighboring M1 monomers extracted from the full reconstruction (grey isosurface), fitted with the atomic model built based on this density. **f)** The model of an individual monomer.

We next co-expressed the IAV HK68 proteins HA, NA, M1 and M2 to form virus-like particles (VLPs) ^4^ and imaged them as described above for virions. The VLPs were essentially indistinguishable from viral filaments. We determined the structure of the M1 layer from the VLPs (**Extended data Fig. 2d-f**) finding it to be identical to that obtained from virions (**Extended data Fig. 2g-i**), confirming our assignment of the protein density as M1. Due to its more isotropic resolution, we used the M1 reconstruction obtained from VLPs (**Extended data Fig. 2f**) for further interpretation.

At the resolution obtained (~8 Å), alpha helices within individual M1 monomers were clearly observed. The crystal structure of the M1 NTD monomer determined at pH 7 (PDB:1ea4 ^11^), a pH most similar to the interior of flu virions, could be fitted as a rigid-body into the membrane proximal lobe of our structure; all nine alpha helices of the M1 NTD monomers are accommodated by the density (**Fig. 2a,b**). Previous biochemical studies have suggested that the M1 NTD interacts with negatively charged phospholipids via electrostatic interactions ^11,21^. Consistent with these observations, in our structure the interaction between M1 and the membrane is mediated by a positively charged surface formed by helices 5, 6 and 8 (**Fig. 2c**).

Fitting multiple copies of the M1 NTD structure reveals that M1 polymerizes into parallel linear strands (**Fig. 2a**). Within the strands, loops between NTD helices form an interface which resembles the crystal packing in the M1 NTD crystal obtained at neutral pH (1ea3,^11^) (**Extended data Fig. 3c**). Parallel M1 strands in the virus are packed tightly together. Each M1 NTD appears in close proximity to two monomers on each adjacent strand (**Extended data Fig. 3a**). Positively charged residues from helices 6 and 7 face negatively charged residues in the parallel strand, contributed from helix 3 of one monomer and the loop between helix 4/5 in another (**Extended data Fig. 3b**). The interface between monomers from adjacent strands is small (~360 Å^2^), suggesting a weak interaction that may allow strands to slide relative to each other. Several mutations or variations altering virion morphology ^22–24^ are located at the M1 NTD-NTD inter-strand interface (**Fig. 2c**), suggesting that strand packing may modulate virion morphology.

The density for helix 9 extends beyond the crystallized helix 9, protruding downwards into the inner lobe density which corresponds to the M1 CTD (**Fig. 2c**). The CTD contains 3 cylindrical densities suggesting that it folds into a 3-helix globular domain, and its orientation suggests formation of a *trans* interface in which it interacts with the membrane-distal surface of the NTD from the neighboring M1 monomer in the linear polymer (**Fig. 2b**). A trans-interface between NTD and CTD is present within the crystal packing of the ISAV matrix protein (**Extended data Fig. 3d**) ^13^.

In order to obtain a higher resolution structure of M1, we expressed and purified recombinant full-length M1. We characterized the purified protein by NMR (**Extended data Fig. 4a**), and found that it is monomeric in solution at neutral pH with a radius of hydration (R_H_) of ~25 Å (**Extended data Fig. 4b**). The NTD adopts the same nine-helix fold as present in the previously determined crystal structure. The CTD, in contrast, is largely disordered, consistent with previous observations ^25^ and does not adopt detectable tertiary structure when pH is increased (**Extended data Fig. 4c**). We identified three stretches with some helical propensity, corresponding to residues 171-190 (helix 10), 196-218 (helix 11) and 231-246 (helix 12) (**Extended data Fig. 4d**).

We assembled purified M1 into helical tubes in the presence of nucleic acid and imaged them by cryo-EM (**Extended data Fig. 5**) - similar coiled structures have been observed in flu virions with disrupted M1 layers ^20^. After classification, the best particles were used to reconstruct a map at 3.8 Å resolution, resolving both the NTD and CTD (**Extended data Fig. 5e,f**). The helix exhibits a pitch of 100.0 Å with 32.4 M1 dimers in each helical turn (**Fig. 2d**). M1 dimerizes via its NTD membrane-binding surface giving rise to two antiparallel linear polymer strands, introducing a D1 symmetry and making the helix apolar. Two nucleic acid filaments are found inside the helix, one at the NTD-NTD dimerization interface and the other in a groove formed at the CTD-CTD interface (**Fig. 2d**, **Extended data Fig. 6c**) – both appear to be nonspecifically bound and only unfeatured density was observed. The 3.8 Å resolution density map allowed building of an atomic model for the full-length M1 protein (**Fig. 2e,f**, **Fig. 3**).

**Figure 3:**
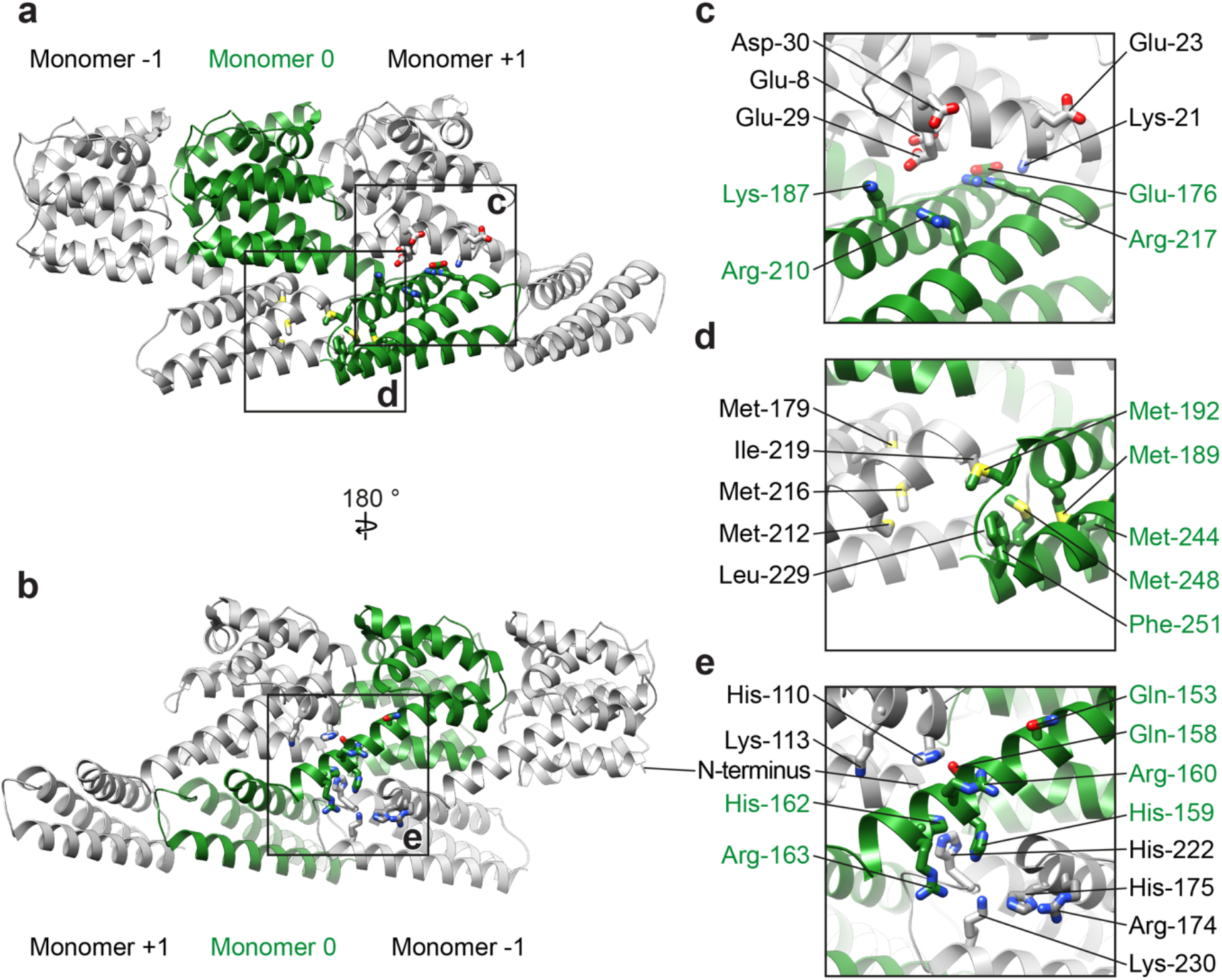
Interactions mediating assembly of M1. **a)** Three neighboring M1 monomers from an M1 strand viewed as in Fig. 2e. **b)** A 180° rotated view of a). In a) and b), residues involved in monomer polymerization are shown in stick representation. Boxes indicate regions magnified in c)-d). **c)** Charged residues mediate interactions between the CTD of monomer 0 (green) and the NTD of monomer +1. **d)** Hydrophobic residues mediate interactions between neighboring CTDs. **e)** A histidine-rich, positively-charged cluster is formed at the interface of three M1 monomers.

The linear M1 strand from the in vitro helical assembly curves in a direction orthogonal to that of the M1 strand from the virus, and with opposite handedness. The structure and arrangement of M1 within the linear M1 strands is, however, very similar (**Fig. 2**, **Extended data Fig. 6**). In both cases the same surfaces of the NTD mediate polymerization, placing the C-terminus in the same trans-interacting position. We therefore interpret the in vitro assembly as a higher-resolution model for the structure and arrangement of M1 strands within the virion.

The structure of the NTD is essentially identical to that of N-terminal fragment structures previously determined by X-ray crystallography (1aa7 ^10^,1ea3 ^11^, 5v6g ^26^) with the exception of the helix4/5 loop which adopts a different conformation (**Extended data Fig. 6).**

The last helix in the NTD, helix 9, extends by three helical turns beyond the crystal structure, before proline-171 bends the peptide chain backwards to form the CTD. The CTD is composed of 82 residues organized into 3 helices (**Fig. 3**) found at the positions where helical propensity was highest in the NMR experiments (**Extended data Fig. 4**). The extension of helix 9 positions the CTD below the membrane distal surface of the neighboring NTD, as in the viral structure, where it forms a tight interaction (**Fig. 3a-c**). The NTD-CTD interface appears to be stabilized by charge complementarity between the negatively charged membrane-distal surface of the NTD (residues Glu-8, Glu-23, Glu-29, Asp-30), and the positively charged interacting surface of the CTD (residues Lys-187, Arg-210, Arg-217), as well as by an inter-molecular salt bridge between Lys-21 and Glu-176. (**Fig. 3c**).

The three-helix bundle of the CTD is stabilized by two methionine-rich hydrophobic cores, one containing Met-189, Met-192, Met-244, Met-248 and the other containing Met-179, Met-212, Met-216 (**Fig. 3d**). The two hydrophobic cores from neighboring monomers in the strand interact with one another via a hydrophobic interface including Ile-219, Leu-229 of one monomer, and Met-192, Met-248, Phe-251 of the neighboring monomer (**Fig. 3d**).

Overall, linear polymerization of full-length M1 by trans-interactions between the NTD and the neighboring CTD buries a surface area of 2137 A^2^ per monomer, compared to 776 A^2^ contributed by the NTD (1-158) alone. These extra interfaces contribute considerable energy to promote M1 polymerization and virion filament extension.

The helix 9 extension interacts with neighboring M1 monomers in both the +1 and −1 positions (**Fig. 3a,b,e**). Directly adjacent to the helix 9 extension we resolve an interaction between the N-terminus of the +1 M1, and the loop between helices 11 and 12 of the −1 M1 (**Fig. 3e**): there are therefore direct interactions between three M1 monomers centred around the helix 9 extension. All three monomers contribute to form a highly positively charged amino acid cluster of 5 arginine or lysine residues and 5 histidine residues. The helix 9 extension of the central monomer contributes His-159, Arg-160, His-162, and Arg-163; helix 7 in the NTD of the +1 M1 monomer contributes His-110 and Lys-113; the CTD of the −1 M1 monomer contributes residues Arg-174, His-175, His-222, and Lys-230 (**Fig. 3e**). A sequence analysis of M1 from 5600 full IAV genomes from NCBI reveals that this cluster is highly conserved suggesting this feature is important for the influenza lifecycle. Indeed four of the five histidines are strictly conserved in IAV (**Extended data Fig. 7**), and while His-222 to glutamine substitutions are found in certain H9N2 viruses ^27^ and His-159 to tyrosine in some bat derived H17N10 and H18N11 sequences, they are compensated by substitution of Gln-158 to His, and Gln-153 to His, respectively (**Fig. 3e**, **Extended data Fig. 7**). Histidine is known to function as a pH sensor and histidine clusters have been described to regulate structural changes in response to low pH in diverse viruses ^28,29^. The M1 layer disintegrates and dissociates from the inner membrane leaflet when influenza virions are exposed to low pH ^9,30,31^. The protonation of the five histidines in this cluster would introduce additional positive charges, providing a mechanism by which destabilization and disassembly of the M1 layer can be triggered in the endosome.

Our observations show that IAV M1 polymerizes into parallel linear strands which assemble helical arrays to form the viral endoskeleton. Polymerization is associated with extension of helix 9 and transition of the CTD from an unfolded to folded state, upon which it interacts with the NTD of the neighboring M1 molecule in the strand. This arrangement implies that virus assembly will be processive – binding of M1 into the filament induces conformational changes that create new interfaces promoting binding of the next monomer. Assembly buries a large surface area, providing an energy source to drive assembly and protrusion of the virion. Once assembled, five histidines that are distant in sequence and contributed by three sequential M1 monomers in the filament come together to form a histidine cluster that can serve as the trigger for pH-mediated M1 disassembly.

## Methods

### Cell lines

Human embryonic kidney 293-T (HEK293T) and Madin-Darby canine kidney (MDCK) cells were cultured in cell culture medium: Dulbecco’s modified Eagle medium (DMEM) Glutamax (Gibco) supplemented with 10% Fetal Bovine Serum (FBS) and 1% penicillin/streptavidin (PS).

### Preparation of HK68 virus samples

Initial virus stocks of A/Hong Kong/1/1968 (H3N2) (HK68) influenza A virus (IAV) generated by reverse genetics as described in ^32^ were amplified in MDCK cells ^33^. All experiments with HK68 virus were carried out under BSL-2 conditions. In preparation for cell seeding, QF AU-200 mesh R2/2 grids (Quantifoil) were glow-discharged and placed into a 6-well tissue culture plate well containing 2 mL of cell culture medium for at least 1 h before cell seeding. 150 000 MDCK cells diluted in cell culture medium were seeded per well and incubated at 37 °C 5% CO_2_, 100% humidity for 12 h to 24 h until cells were adhered to EM grids. Wells were washed with 0.3% BSA/10 mM HEPES in DMEM before inoculation with diluted virus at an multiplicity of infection (MOI) of 1. Cells were incubated with virus for 1 h, gently shaking every 15 min. After 1 h, the inoculum was removed and replaced with 0.3 % BSA/DMEM containing 0.5 μg/ml to 5 μg/ml TPCK-treated trypsin, depending on trypsin activity. Infection was stopped when cytopathic effect (CPE) was visible by light microscopy, typically 24 h to 48 h post infection. Grids were removed from cell culture medium, and 5 μL of 1:3 diluted 10 nm colloidal gold in PBS was added to each grids prior to plunge-freezing by back-side blotting using a LeicaGP cryo plunger (Leica).

### Preparation of HK68 HANAM1M2 VLP samples

The sequences of A/Hong Kong/1/1968 (H3N2) (HK68) HA, HK68 NA, HK68 M1 and HK68 M2, cloned into pCAGGS expression vectors as previously described in ^4^, were used for the production of HK68 VLP samples. VLP samples were prepared equivalently to HK68 virus samples. 300,000 HEK293-T cells were seeded per well with grids present inside the well. When cells were adhered to EM grids, typically 12-24 h post seeding, cells were co-transfected with the four plasmids in the ratio 1:1:2:0.5 for HA, NA, M1, M2, using the transfection reagent FUGENE (Promega) according to the manufacturer’s protocol. 48 h post-transfection, grids were removed from cell culture medium and plunge-frozen as described above.

### Cryo electron tomography

Collection of cryoET data was performed as described previously ^34^. Tilt series were acquired on a Titan Krios (Thermo Fisher, formerly FEI) operated at 300 keV equipped with a Gatan Quantum 967 LS energy filter using a 20 eV slit in zero-loss mode. Images were recorded on a Gatan K2-Summit direct electron detector with a pixel size of 1.78 Å or 1.7 Å at the specimen level. Tomograms were acquired from −60° to 60° with 3° increment and a dose-symmetric tilt-scheme ^35^ using SerialEM-3.49 ^36^, with defoci between −2 μm and −6 μm, 20 Frames were aligned using the alignframes function in the IMOD software package ^37^. If data were collected in super resolution mode, frames were Fourier cropped to 4K x 4K pixel images. The contrast transfer functions (CTF) for each tilt image was determined on non-exposure filtered tilt stacks using CTFFIND4 ^38^ and evaluated for each image. Tilt images were exposure filtered according to the accumulated dose applied to the sample for each tilt as described in ^39^ using Matlab (MathWorks) or the dose weight filtering options in alignframes function in IMOD 4.10. Tilt series were aligned in etomo/IMOD using gold fiducials. Tomograms with less than three trackable fiducials were discarded. Tilt images were CTF-multiplied and reconstructed by weighted back-projection using NovaCTF ^40^. Tomograms were subsequently isotropically binned by a factor of two and by a factor of four (referred to as Bin2 and Bin4 tomograms in the following). Detailed imaging parameters per dataset are summarized in **Extended data Table 1**. Tomograms were visualized either in IMOD or UCSF Chimera 1.13.1 ^41^.

### Subtomogram averaging

Tomograms which were acquired at a defocus range from −1 μm to −4 μm were used for subtomogram averaging. Data from HK68 virus tomograms and HK68 HANAM1M2 VLP tomograms were processed using essentially the same strategy.

Subtomogram averaging was performed using scripts in Matlab (Mathworks) based on functions from the AV3 ^42^, TOM ^43^ and Dynamo ^44^ software packages. To determine initial positions and orientations for subtomogram extraction, the central axis of each virus/VLP filament was marked in Bin4 tomograms using the Volume Tracer function in UCSF Chimera and fitted with a spline function. A tubular grid of points was generated at the radial position of the matrix layer, centered on this spline, with a grid spacing of 2 Bin4 pixels (approximately 1.5 nm). This generated 20,000 - 35,000 positions per filament. Overlapping cubic subtomograms with box size 64 pixels (46 nm) were extracted at these grid positions. Initial Euler angles were assigned based on the orientation of the normal vectors relative to the tube surface at each grid position. For HK68 virus data, subtomograms were split into two independent data half sets, based on their location in either the upper or the lower part of each filament, prior to starting any subtomogram averaging or alignment.

Subtomograms for each filament were averaged to generate starting references, and then iteratively, rotationally and translationally aligned and averaged at Bin4, and then at Bin2 (in 96 pixel boxes) to generate low resolution structures of the M1 layer for each filament. Structures from individual filaments differed in radius, number of helix starts, and radius (**Fig.1**, **Extended data Fig. 1**). M1 subtomograms extracted from 4 filaments with the same parameters (3-start, right-handed M1 layers) were combined. For VLP data, subtomograms were divided into two half sets, based on location in either the upper or the lower part of each filament, after assignment of helix start number, and further processing was performed independently for the two half sets. Subtomograms were re-extracted at Bin1 (in 120 pixel boxes) and final iterations of alignment were performed applying a bandpass filter from 8 Å to 35 Å. The NTD and CTD layers were subsequently refined independently using the final alignment parameters and masks which focused on the respective layer.

For HK68 virus data the missing wedge was modelled at all processing stages as the sum of the amplitude spectra of subtomograms extracted from regions of each tomogram containing empty ice, and was applied during alignment and averaging. For VLP data, this wedge was applied from Bin1, a binary wedge was used at earlier alignment stages.

For each of the refined layers, we assessed the regularity of the local packing and the quality of alignment by plotting the distribution of positions of all neighboring subtomograms for each M1 position in a so-called ‘neighbor plot’ as described in ^45^. The peaks corresponding to the positions of neighbors in the helical lattice were elliptical and extended along the tube surface, perpendicular to the tube axis (the Y-direction). This observation suggests either higher variability in the position of neighbors in this direction, or lower accuracy of alignment in this direction. We selected the subsets of M1 monomers where at least three neighboring M1 monomers were found within defined spheres centered on the peak neighbor position. This step reduced resolution anisotropy while retaining ~50 % of subtomograms. The cleaned, refined maps of the NTD and CTD layers were then recombined into the final structure (**Fig. 1**, **Extended data Fig. 2**). Density maps were visualized either in IMOD or UCSF Chimera.

### Directional resolution measurements

Anisotropy in resolution was assessed by calculating the Fourier shell correlation (FSC) within cones of 40° ^46^. Local and global resolution measurements and the determination of a global B-factor were performed in RELION by FSC between the two independent half-datasets within a smooth-edged mask including the M1 layer. Final structures were sharpened using the determined B-factor and filtered according to the determined global resolution ^47^.

### Structure fitting and modelling based on the M1 in situ structure

The crystal structure of the M1-NTD determined at neutral pH, (PDB:1ea3) ^11^ was fitted as a rigid body into the subtomogram averaging structure using the fit in map functionality in UCSF chimera. The lengths of CTD alpha-helix-like densities were measured in UCSF Chimera and cylinders of the measured length were placed into the density to illustrate the secondary structure of the M1 CTD.

### Expression and purification of M1 for NMR measurements

For NMR, *Escherichia coli* Rosetta 2 (DE3) bacteria (Merck millipore) transformed with the pET21b (Merck millipore) vector carrying the coding sequence of wildtype PR8 M1 were grown in 99.9% ^2^H_2_0 (cortecnet) M9 medium, containing 5.4g/L Na_2_HPO_4_ (anhydrous), 2.7g/L KH_2_PO_4_ (anhydrous), 0.45g/L NaCl, 1.5 g/L yeast nitrogen base (YNB) (Sigma-Aldrich Cat # Y1251), 3.5 g/L ^13^C,^2^H D-Glucose (Sigma-Aldrich Cat # 552151) and 1g/L ^15^NH_4_Cl (Sigma-Aldrich Cat # 299251). Prior to large-scale expression cells were adapted for growth in deuterated media on M9 medium agar plates (1.5% w/v) containing 10, 44 and 78% ^2^H_2_O.

The isotopically labelled M1 protein was purified as stated below for in vitro assembly of M1 helical tubes, in citrate buffer (50 mM citrate pH 5, 50 mM NaCl) and subjected to additional cation exchange chromatography using a HiTrap SP HP column (GE lifesciences) to remove unbound bacterial DNA. The eluted M1 protein was buffer exchanged into 50 mM citrate buffer (50 mM citrate pH 5, 50 mM NaCl), concentrated to 2 mg/mL (~150 μM) and 5% ^2^H_2_O, 0.02% NaN_3_ (Sigma Aldrich) and 0.25 mM 4,4-dimethyl-4-silapentane-1-sulfonic acid (DSS) were added.

### NMR

All NMR experiments on M1 PR8 were carried out at 25°C on a Bruker Avance II+ 700 MHz spectrometer equipped with a triple resonance TCI cryo probe. From 1D experiments the sidechain deuteration level was estimated to be above 90%. For the backbone chemical shift assignment, we recorded non-linear sampled TROSY ^48^ versions of ^1^H-^15^N HSQC (full sampling), HNCO, HNcaCO, HNCA, HNcoCA, HNCACB, HNcoCACB, CCcoNH and hNcocaNNH spectra. The spectra were processed and/or reconstructed using either nmrPipe ^49^, TOPSPIN 4.0.8 or compressed sensing implemented in qMDD (mddnmr) ^50^. Backbone resonance assignments were performed using ccpnmr ^51^. Cα, Cβ and C’ secondary chemical shifts were calculated using random coil chemical shifts provided by ^52^.

The pH titration was performed on a non-deuterated sample. We recorded ^1^H-^15^N HSQCs, and X-STE diffusion experiments at pH points 5, 5.5, 6, 6.5, 7, 7.5, 8, 9 and 10.

^15^N-edited ^1^H heteronuclear stimulated-echo longitudinal encode-decode diffusion experiments (X-STE) incorporating bipolar pulse paired gradients ^53^ were used to measure diffusion coefficients of exclusively ^15^N-labelled species, using a diffusion delay ∂ of 100 ms and 2*2ms gradient pairs *δ* for encoding and decoding respectively. Peak intensities I_o,_ I_j_ at multiple gradient strengths (5-85% of the maximum allowed gradient strength) were integrated and the diffusion coefficient was calculated using Stejskal-Tanner equation, where I_0_ corresponds to the peak intensity observed at the weakest gradient strength and I_j_ the peak intensity at a given gradient strength *G* in Gauss*cm^−1^, with *tau* corresponding to the gradient recovery delay and γ is the ^1^H gyromagnetic ratio:

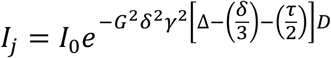

The hydrodynamic radius was calculated according to the Stokes-Einstein equation *R_h_ = kT / (6πηD)* where *k* is the Boltzmann constant, *T* the absolute temperature and *η* the solvent viscosity. Theoretical radii of hydration were calculated in HYDROPRO ^54^ based on the PDB files of the isolated NTD and the full-length protein.

### Protein production and in-vitro assembly of M1 helical tubes

Full-length A/Puerto Rico/8/1934 (H1N1) (PR8) M1 cDNA with Arg134Lys substitution was cloned into pET21b resulting in a C-terminal extension LEHHHHHH. The Arg134Lys substitution was found to improve sample homogeneity during tube assembly and this substitution has been found to occur naturally in M1. Protein was expressed by 0.1 M IPTG induction in Rosetta (DE3) cells at 37 °C for 3 hours in LB media. Cells were pelleted and lysed in Tris-saline (50 mM Tris-HCl pH 8.0, 150 mM NaCl) in the presence of 0.1% Triton X-100, and 0.1 mg/ml egg-lysozyme. The lysate was layered over a 60% sucrose cushion made in tris-saline and supplemented with 0.1% triton X-100 and centrifuged in a SW-32 rotor for 30 min at 25000 RPM. The insoluble pellet from the centrifugation step was found to contain mostly M1 and negative stain EM show it exists as aggregated tubular structures (**Extended data Fig. 5a**). Insoluble M1 pellet was resuspended in citrate-buffer (50 mM citrate pH 4.6, 50 mM NaCl), and after centrifugation at 20000 g for 10 minutes, soluble M1 was located in the supernatant. To assemble M1 into tubes, solubilized M1 was mixed with a purified 6.4 kb size plasmid DNA in a w/w ratio of 1:2.5 and adjusted to an M1 protein concentration of 0.1 mg/ml in glycine-saline (100 mM glycine pH 10, 150 mM NaCl). The tube assembly reaction was carried out at 21 °C for 16 hours and checked by negative stain EM (**Extended data Fig. 5b**).

### In vitro helical tube CryoEM sample preparation

C-Flat 2/2-3C grids (Protochips) were glow discharged for 30 seconds at 25 mA, and a 5 ul sample of the tube assembly reaction was applied 3 times to the grid with 30 s incubation and blotting between each sample application. Grids were washed with 5 μl H_2_O before blotting and plunge-freezing in liquid ethane using an FEI Vitrobot. Grids were stored in liquid nitrogen until imaging. Imaging was performed on an FEI Titan Krios operated at 300kV, equipped with a Gatan K2-summit direct detector using a 20 eV slit width. 2347 2D images were acquired using SerialEM-3.7.0 ^36^ with a nominal magnification of 105000 X, giving a pixel size of 1.128 Å at the specimen level.

### 2D CryoEM image processing

All super-resolution frames were corrected for gain reference, binned by a factor of 2, motion-corrected and dose-weighted using MOTIONCOR2 ^55^. Aligned, non-dose-weighted micrographs were then used to estimate the contrast transfer function (CTF) using CTFFIND4 ^38^.

Filaments were picked manually, and segments were extracted using a box size of 564 Å and an inter-box distance of 33 Å, obtaining 463152 helical segments. Reference-free 2D classification was carried out in RELION 3.0.8. ^56^ 2D classes with clear structural features were selected resulting in 297183 helical segments. Analysis of selected 2D class averages using Spring package ^57^ using segclassexam revealed that tube diameters varied between 315-350 Å. The most abundant classes (those with diameters between 327-329 Å as measured by segclassexam) were selected in RELION resulting in 61058 helical segments. Using one 2D class average with minimal out-of-plane tilt, segclassreconstruct within Spring was used to test possible helical parameters, identifying a pitch of ~100 Å and 27.8, 29.8, 31.8 or 28.2, 30.2, 32.2 subunits per turn. These helical parameters were tested in RELION 3D refinement and 32.2 subunits per turn as an initial helical parameter resulted in a density showing protein features, the refined pitch was 99.8 Å and 32.4 subunit per turn. 3D-classfication using a mask focusing on the central 50% of the M1 helical tube further reduced heterogeneity in diameter – the best 3D class contained 17984 helical segments and was used for final 3D auto-refinement and Bayesian polishing in RELION ^58^.

Overall resolution estimates were calculated from Fourier shell correlations at 0.143 criterion between two independently refined half-maps, using phase-randomization to correct for any convolution effects of the generous, soft-edged solvent mask focusing on the central 25% of the helix, that is ~1.4 helical turns. Final reconstructions were sharpened using standard post-processing procedures in RELION, resulting in a *B*-factor of −115 Å^2^ (**Extended data Table 2**). Helical symmetry was imposed on the post-processed maps using the relion_helix_toolbox ^59^.

### Model building and refinement

The N-terminal fragment crystal structure (PDB:1AA7 ^10^) was fitted into the sharpened 3.8 Å map and residues were manually adjusted, C-terminal domain residues were built manually in Coot-0.9 ^60^. The final model contains residue 2-252, the N-terminal methionine is removed by cellular methionine aminopeptidase while the density for the C-terminal tag is present but poorly defined for model building. The structure was refined in PHENIX-dev-3580 ^61^ to good geometry with no Ramachandran outliers and statistics are given in (**Extended data Table 2**).

## Data availability

The cryo-EM and cryo-ET structures, and representative tomograms are deposited in the Electron Microscopy Data Bank (EMDB) under accession codes EMD-XXXXX - EMD-XXXXX. The associated molecular model is deposited in the protein data bank (PDB) under accession XXXX.

## Acknowledgments

We thank Oliver Schraidt, Susann Kummer, Vera Sonntag-Buck, Vojtech Zila, Petr Chlanda, Steffen Klein, Dustin Morado, Sjors Scheres, Conny Yu, Cheng-Yu Huang and Wim Hagen for technical assistance, and Peter Rosenthal, Steve Gamblin, John Skehel and David Veesler for support during preparation of helical arrays. All NMR data were acquired at the MRS facility of the MRC-LMB. This study made use of electron microscopes at EMBL and the MRC-LMB EM Facility, as well as high-performance computing resources at EMBL and LMB and the CIID CL2 imaging facility, we thank the staff who maintain those resources. Funding was provided to JAGB by the European Research Council (ERC) under the European Union’s Horizon 2020 research and innovation programme (ERC-CoG-648432 MEMBRANEFUSION), the Medical Research Council (MC_UP_1201/16) and the European Molecular Biology Laboratory; and to JAGB and HGK by the Deutsche Forschungsgemeinschaft (project number 240245660 - SFB1129).

## Author contributions

XX, HGK and JAGB conceived the project. XX designed and performed in vitro helical assembly of M1, LJC contributed to initial identification of helical arrays. SE and SF designed and performed NMR experiments. JP and JAGB designed virus and VLP experiments which were performed by JP. JP, XX, and KQ collected cryo-EM and cryo-ET data. JP processed cryo-ET data with assistance from WW and JAGB. XX processed cryo-EM data with assistance from KQ and JAGB. JP, XX, SE and JAGB interpreted data. JP, XX and JAGB wrote the original draft which was edited and reviewed by all authors. JAGB and HGK obtained funding. JAGB supervised and managed the project.

## Extended data

**Extended Data Figure 1:**
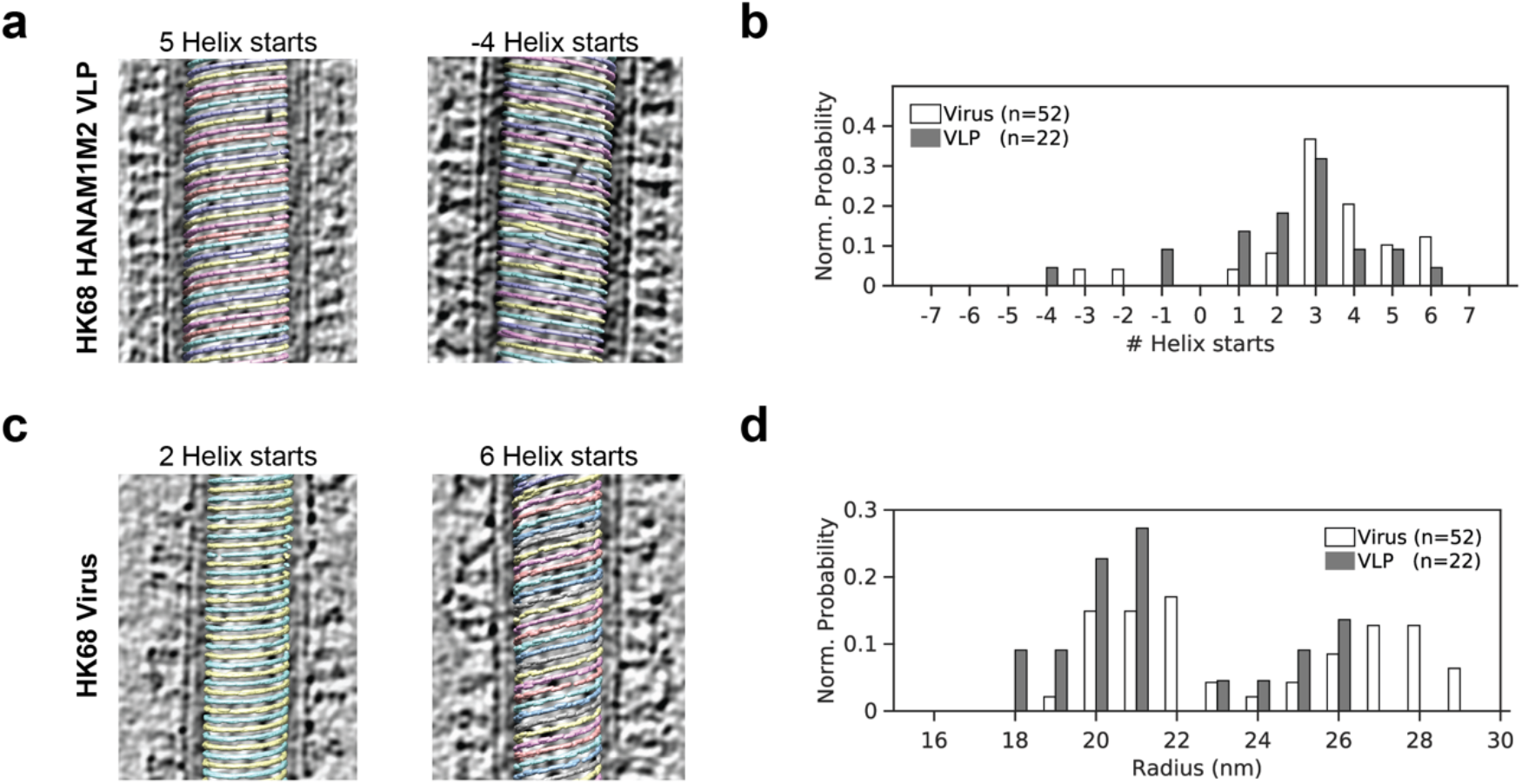
HK68 virions and VLPs have variable numbers of M1 strands and variable radius. **a)** Slices through tomograms of two HK68 VLP filaments, superimposed with a visualization of M1 subtomogram positions and orientations. Separate M1 strands are shown in different colors. Left: M1 arranged as 5 parallel right-handed helical strands and Right: 4 parallel left-handed helix strands. **b)** Histogram showing the distribution of number of helical strands (# Helix starts) for all HK68 virus (white) and VLP (grey) filaments analyzed. Right-handed helices have positive start numbers, left-handed helices have negative start numbers. **c)** As in A) but for two HK68 virus filaments. **d)** Histogram showing the distribution of filament radius for all HK68 virus and VLP filaments analyzed. Radii were determined at the position of the M1 NTD.

**Extended Data Figure 2:**
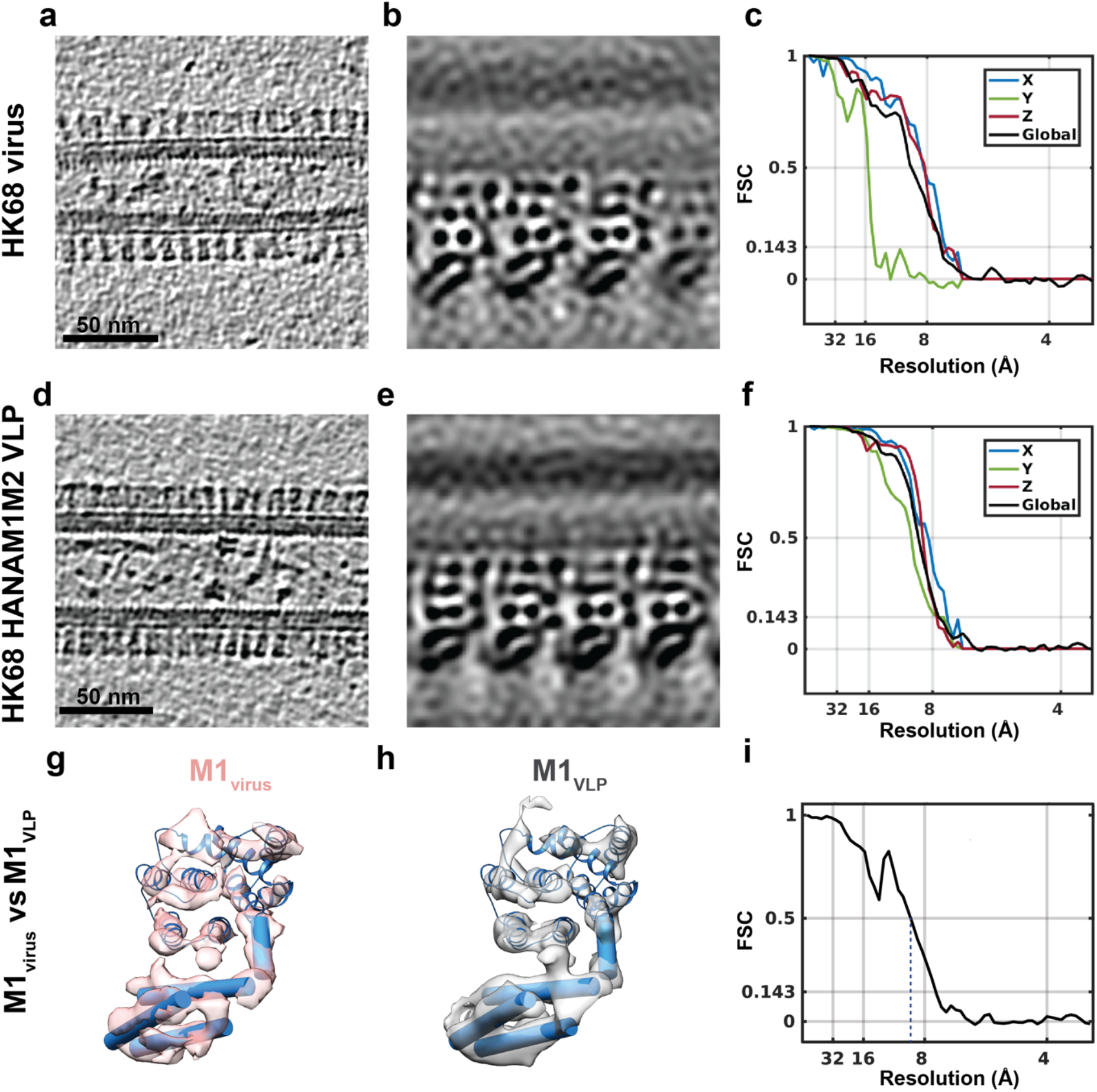
Comparison of HK68 virus and VLP M1 structures, and resolution measurements. **a)** Representative orthoslice through a HK68 virus tomogram. **b**) Projection through 1.8 nm of the M1 subtomogram average obtained from HK68 virus data. **c**) Global FSC curve for the structure of M1 from HK68 virus (black), and analysis of anisotropy by FSC curves in X, Y, and Z directions (blue, green red). **d**) - **f**) As in **a**) - **c**) for VLP data. **g**) The structure of M1 from HK68 virus derived from subtomogram averaging, shown as a pink isosurface and fitted with the M1 NTD crystal structure. Cylinders represent the secondary structure elements of the M1 CTD. **h**) As in **g**) but for the subtomogram averaging structure of M1 from VLPs. **i**) FSC calculated between the virus M1 and VLP M1 structures indicating that the structures are the same up to resolution of 9 Å.

**Extended Data Figure 3:**
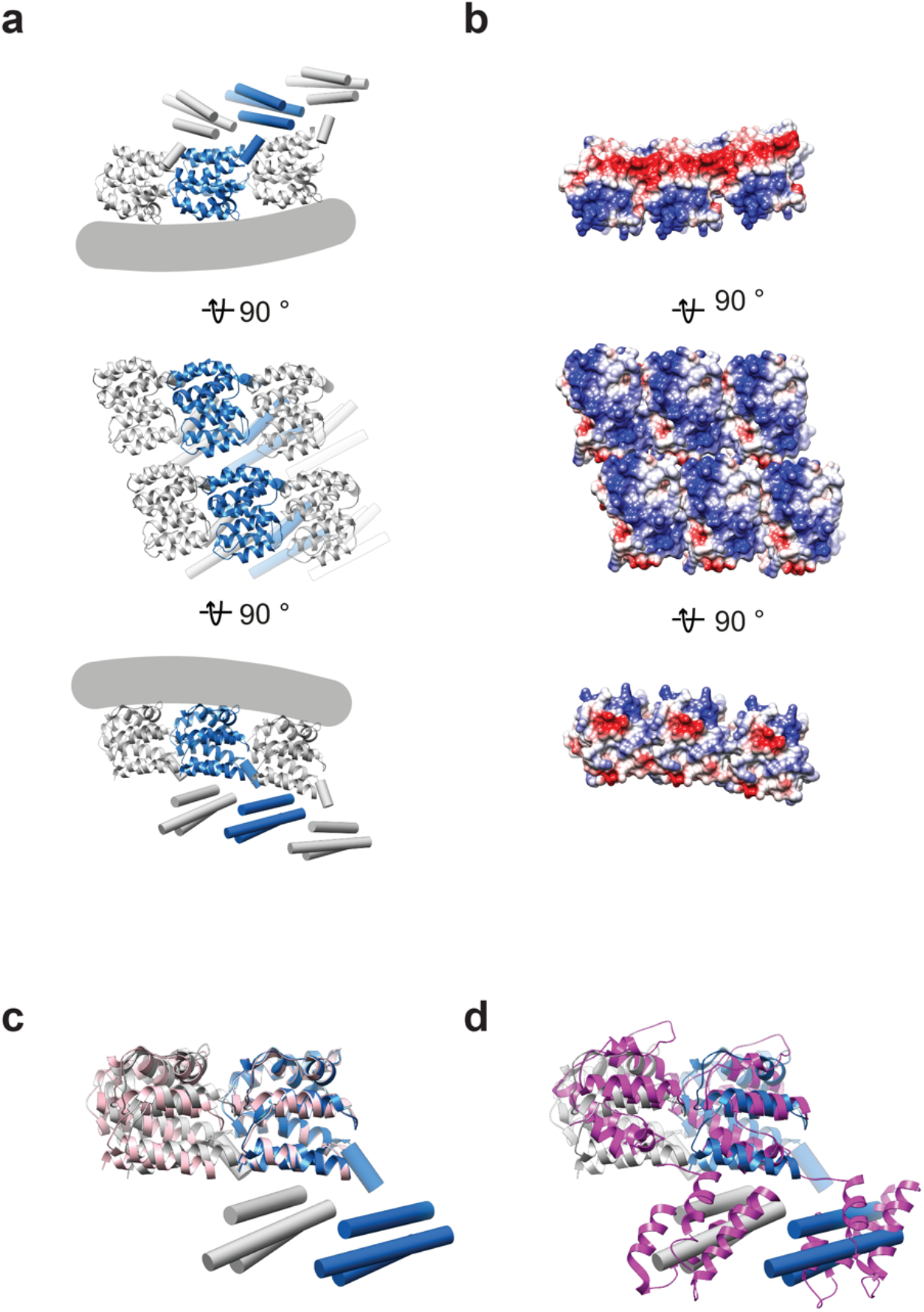
Analysis of the M1 structure determined within virions. **a)** Model of M1 subunits as they are arranged in the virus, shown in three different orientations (as in Fig. 2b.) **b)** Surface charge representation for the NTD in equivalent orientations to those shown in a). **c)** Overlay of the in-virus M1 model shown in a) with two neighboring M1 NTD domains (pink) within the crystal packing in PDB:1ea3 ^11^. **d)** Overlay of the in-virus M1 model with two neighboring infectious salmon anemia virus (ISAV) matrix proteins (magenta) from the crystal packing in PDB:5wco ^62^.

**Extended Data Figure 4:**
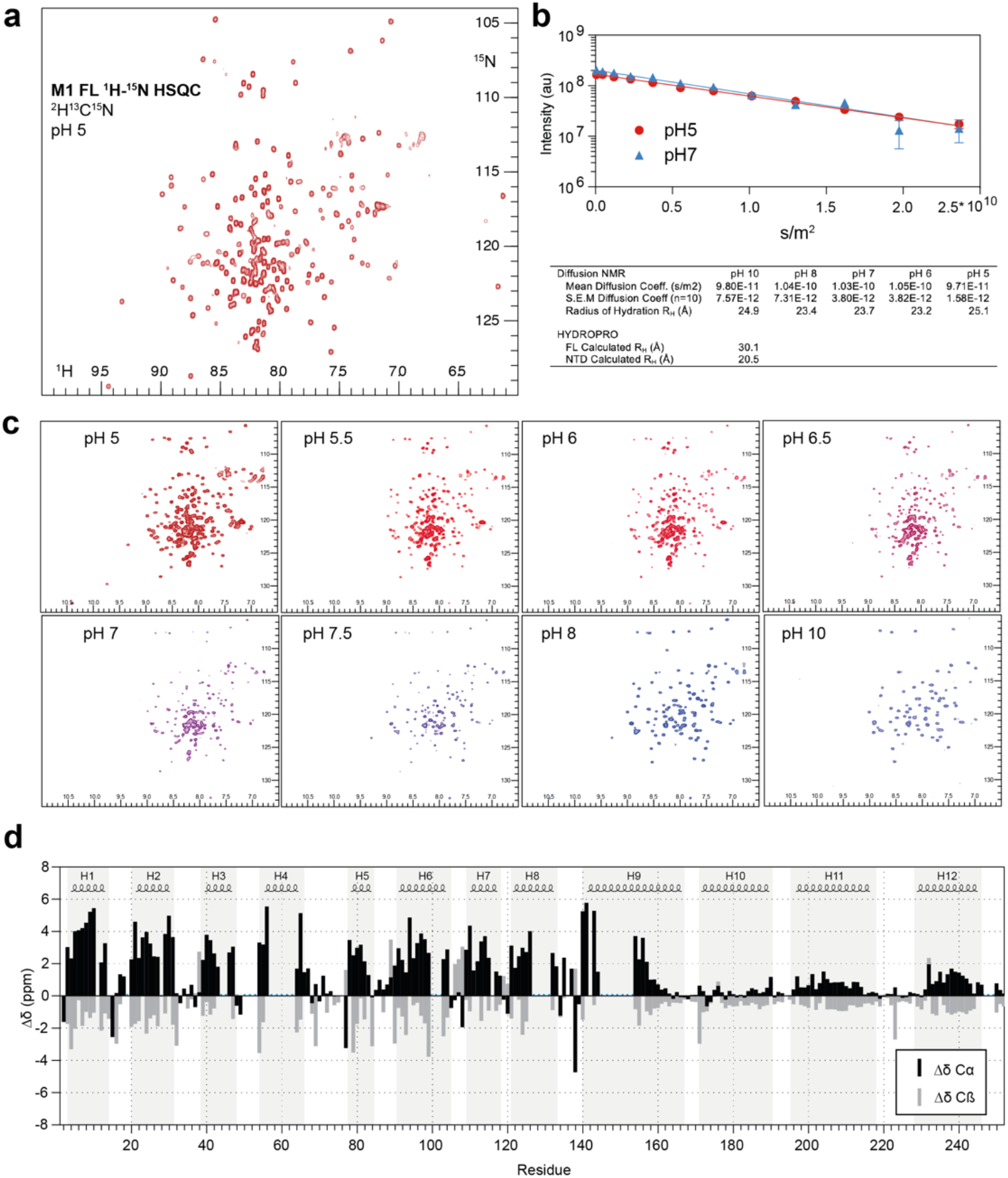
NMR analysis of M1. **a)**^1^H-^15^N HSQC spectrum of full length M1 at pH 5. **b)** Representative diffusion curves acquired at pH 5 and 7 and fitted diffusion coefficients and calculated radii of hydration at pH values 5, 6, 7, 8 and 10. HYDROPRO^54^ calculated radii of hydration are listed below for comparison. **c)** ^1^H-^15^N HSQC spectra of full length M1 at different pH values. Increasing pH in absence of DNA and membrane does not induce folding of the CTD. At high pH (>8) only resonances from the first part of the NTD are visible. The protein remains largely monomeric throughout the pH titration. **d)** Secondary chemical shifts analysis. When compared to random coil, positive Cα and negative Cβ chemical shifts designate helical secondary structure. Helical segments from the single particle structure are depicted above for comparison. Missing assignment are indicated by blue dots.

**Extended Data Figure 5:**
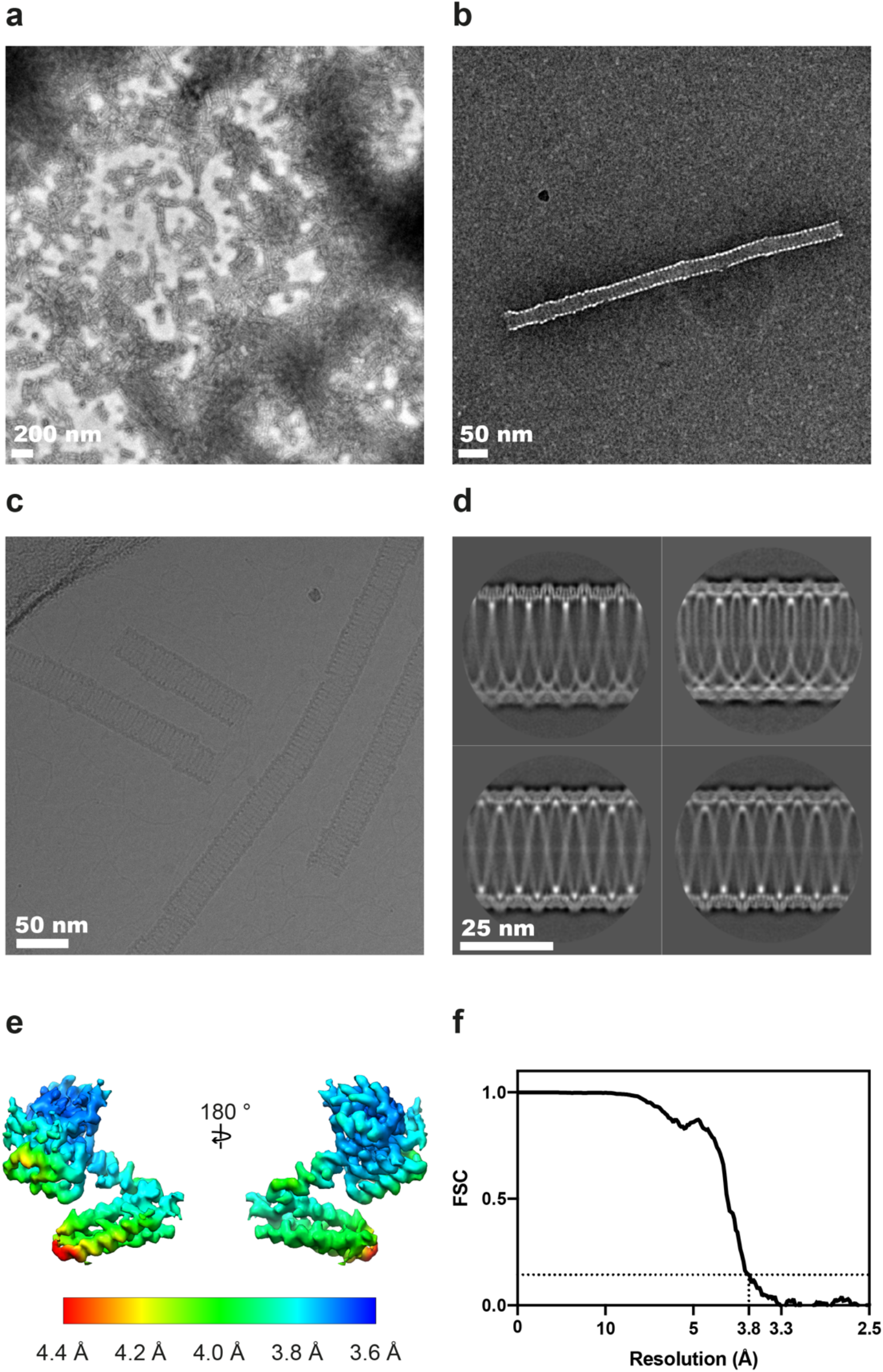
Electron microscopy of *in vitro* reconstituted M1 helical tubes. **a)** Negative-stain image of aggregated M1 tubes found within the pellet after sucrose cushion centrifugation. **b)** Negative-stain image of *in-vitro* assembled M1 tubes in the presence of nucleic acid. **c)** A typical cryoEM image of *in-vitro* assembled M1 tubes at −2.7 μm defocus. **d)** Selected class averages with tube diameter between 327-329 Å as determined by segclassexam. The lower left class average has minimal out-of-plane tilt and was used to test possible helical parameters by segclassreconstruct. **e)** cryoEM density of an M1 monomer is shown, the surface is colored by local resolution of the map as determined by RELION. **f)**Global FSC curve of the final *in-vitro* assembled M1 tube helical reconstruction (**Fig. 2d**).

**Extended Data Figure 6:**
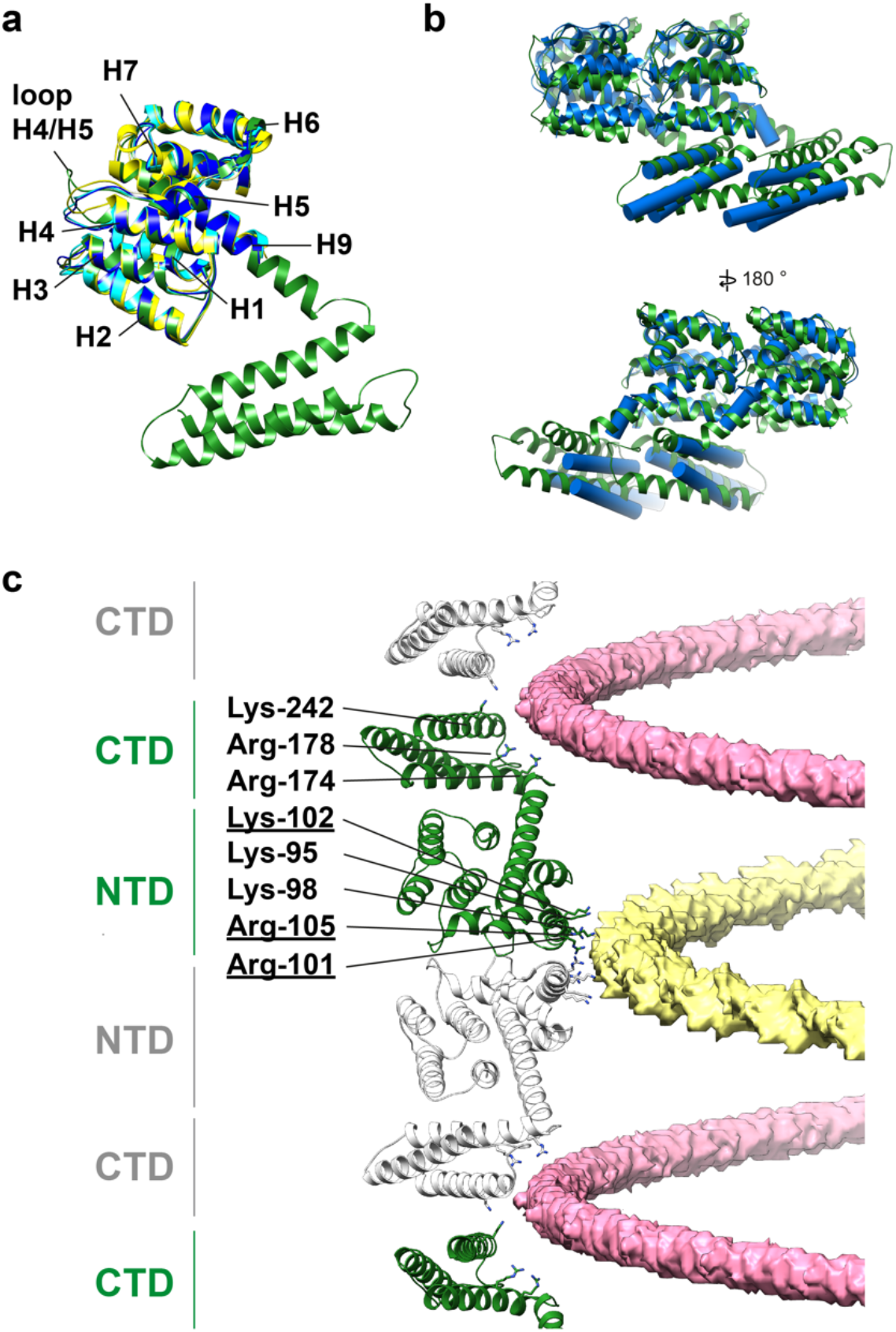
Analysis of the in vitro M1 structure. **a)** Alignment of the full-length M1 structure determined by helical reconstruction (green) to crystal structures of M1 NTD – 1aa7 (blue), 1ea3 (yellow), and 5v6g (cyan). The structures are the same except for small differences in the H4/H5 loop. **b)** Alignment of *in-situ* (blue) and *in-vitro* (green) M1 dimers extracted from their respective linear polymers. The differences are limited to small movements at the interfaces, perturbations in the orientation of helix 9 that accommodate the different curvatures, and a small change in the orientation of helix 12. **c)** A view of M1 monomers extracted from the helical reconstruction to show the sites of interaction with the two nucleic acid strands (see also Fig. 2). One nucleic acid strand binds at the NTD-NTD interface (yellow), the other binds to a groove formed at the CTD-CTD interface (pink). Residues interacting with nucleic acid are all positively charged and are shown as sticks. Residues that form part of the nuclear localisation signal which was previously shown to bind the viral ribonucleoprotein ^63^ are underlined.

**Extended Data Figure 7:**
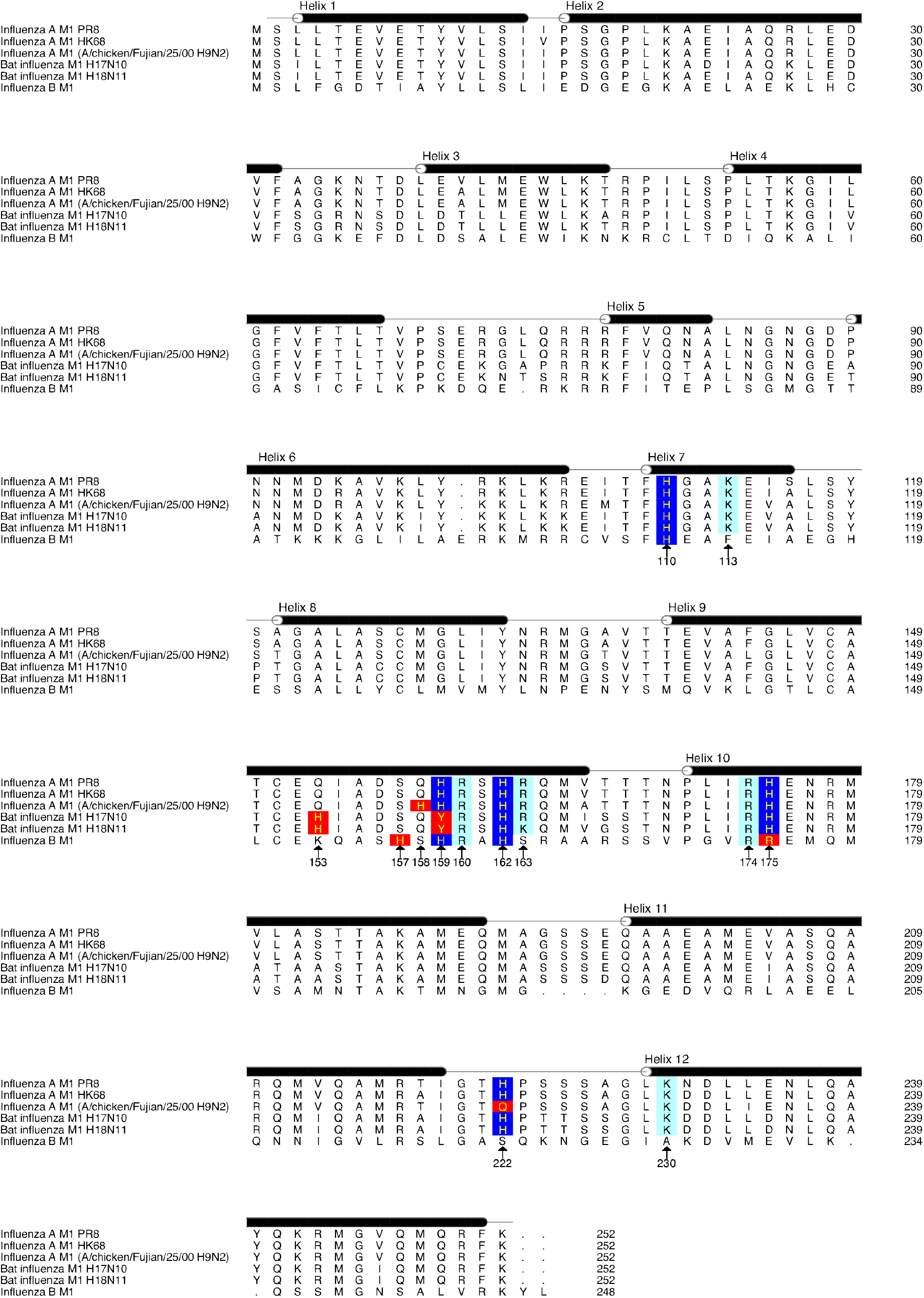
Alignment of M1 protein sequences. M1 sequences of the following viruses: Influenza A M1 PR8 (A/Puerto Rico/8/1934 (H1N1)), Influenza A M1 HK68 (A/Hong Kong/1/1968 (H3N2)), Influenza A M1 (A/chicken/Fujian/25/2000 (H9N2)), Bat influenza M1 H17N10 (A/little yellow-shouldered bat/Guatemala/164/2009 (H17N10)), Bat influenza M1 H18N11 (A/flat-faced bat/Peru/033/2010 (H18N11)), Influenza B M1 (B/Lee/1940) were downloaded from UniProt and aligned using mafft (https://mafft.cbrc.jp/alignment/software/). Locations of Helices 1-12 are marked above the amino-acid sequences. Conserved histidines are shaded blue, conserved charged residues are shaded cyan. Substituted histidine locations and compensatory histidine substitutions are shaded red.

---

**Extended data Table 1.** Data collection and processing parameters for M1 within virions and VLPs.

|  | Influenza A/HK68 virus | Influenza A/HK68<br>HANAM1M2 VLP |
| --- | --- | --- |
| *Data collection and processing* |  |  |
| Magnification | 81,000 X | 81,000 X |
| Voltage (keV) | 300 | 300 |
| Detector | Gatan Quantum K2 | Gatan Quantum K2 |
| Energy-filter | Yes | Yes |
| Slit width (eV) | 20 | 20 |
| Pixel size (Å) | 1.70 | 1.78 |
| Defocus range (µm) | -1.5 to -4.5 | -2 to -6 |
| Defocus step (µm) | 0.25 | 0.25 |
| Tilt range (min/max, step) | -60/60°, 3° | -60/60°, 3° |
| Tilt scheme | Dose-symmetrical<br>(Hagen scheme) | Dose-symmetrical<br>(Hagen scheme) |
| Total Dose (electrons/Å<sup>2</sup>) | ~130-150 | ~120 |
| Frame number | 20 | 20 |
| Tomograms used for M1/acquired | 53/105 | 20/60 |
| Tomograms/viruses used for final structure | 5 | 4 |
| Final subtomograms set A NTD/CTD | 6025/5934 | 6936/7164 |
| Final subtomograms set B NTD/CTD | 6149/6147 | 7611/7822 |
| Symmetry imposed | none | none |
| B-factor | -620 | -665 |
| Map resolution isotropy (Å) at<br>0.143 FSC threshold | 5.8-15 | 6.8– 9.8 |
| Global resolution (Å) at 0.143 FSC threshold | 7.6 | 8.2 |

**Extended data Table 2.**
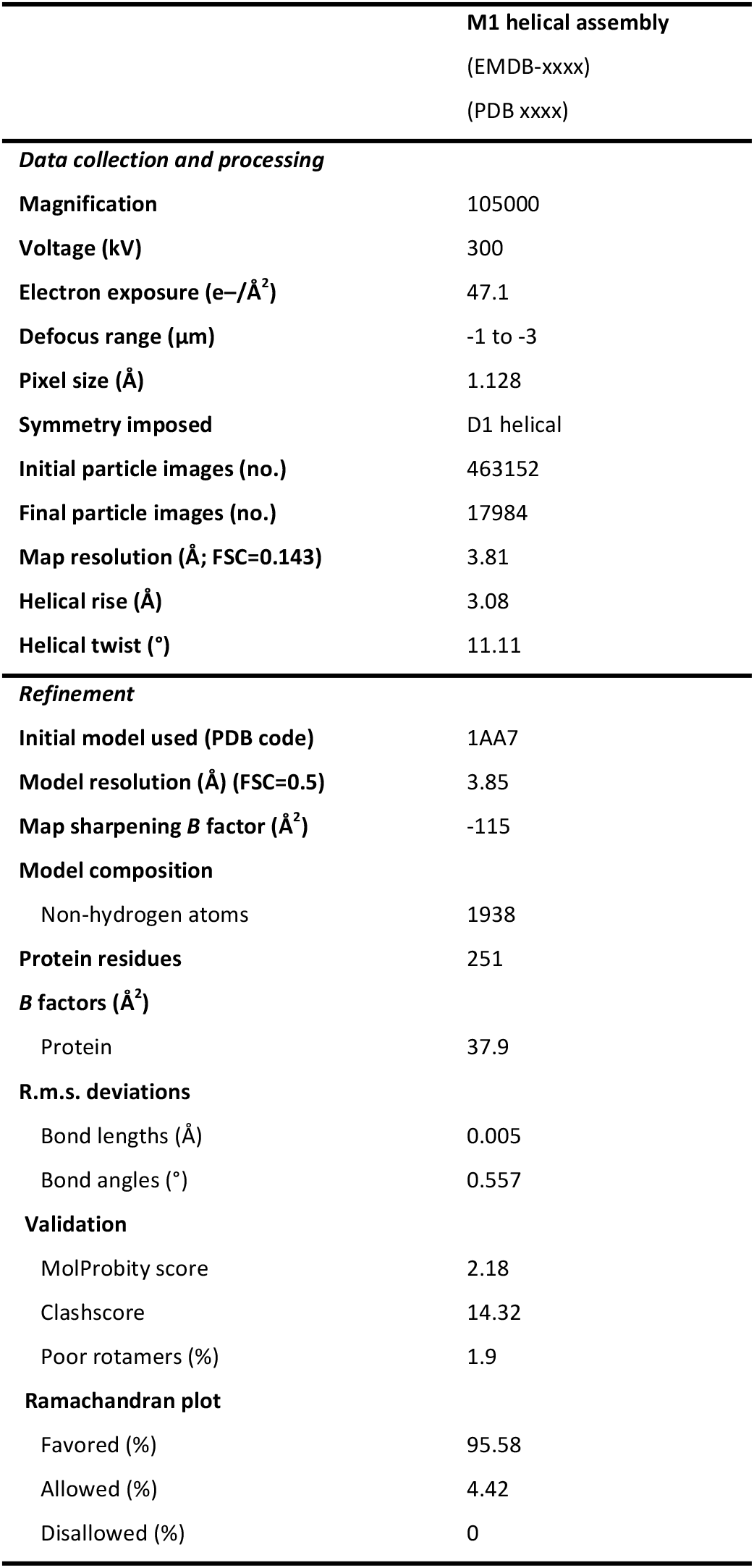
Data collection and processing parameters for in vitro helical assembly of M1.

